# The trichloroethylene metabolite S-(1,2-dichlorovinyl)-l-cysteine inhibits lipopolysaccharide-induced inflammation transcriptomic pathways and cytokine secretion in a macrophage cell model

**DOI:** 10.1101/2022.03.15.484241

**Authors:** Sean M. Harris, Kelly M. Bakulski, John Dou, Ethan Houskamp, Eleanor C. Scheeres, Emily Schellenboom, Olivia Harlow, Rita Loch-Caruso, Erica Boldenow

## Abstract

Previous studies have shown that the trichloroethylene metabolite S-(1,2-dichlorovinyl)-l-cysteine (DCVC) inhibits cytokine secretion in pathogen stimulated fetal membrane tissue but little is known about the mechanism for these effects, including which cell types or transcriptomic pathways are impacted. Macrophages play a critical role in the fetal membrane innate immune response during infection. We tested the hypothesis that DCVC inhibits lipopolysaccharide (LPS) stimulated inflammation pathways in differentiated (macrophage-like) THP-1 cells. THP-1 cells were differentiated with phorbol 12-myristate 13-acetone for 24 hours and subsequently treated with 1, 5, or 10 µM DCVC for 24 hours. After an additional 4 hour incubation with lipopolysaccharide (LPS), we collected RNA and cell media. We performed transcriptomic analysis using RNA sequencing analysis for 5µM DCVC treatments and quantified cytokine release (IL-1β, IL-6, and TNF-α) into cell media for 1, 5 and 10 µM DCVC treatments. RNAseq analysis revealed 1,399 differentially expressed genes (FDR<0.05 and log_2_fold change magnitude>2.5) in the cells co-treated with DCVC and LPS compared to LPS alone. For example, TNF was 9-fold downregulated with the addition of DCVC. Major pathways downregulated (adjusted p-value<0.05) in DCVC+LPS treatments versus LPS-only treatments, included: “acute inflammatory response”, “production of molecular mediator of immune response” and “phagocytosis”. LPS increased IL-1β, IL-6, and TNF-α levels in culture media (p<0.001), but this effect which was inhibited by co-treatment with DCVC (p<0.001 for LPS vs. LPS+DCVC treatments). Our results demonstrate that DCVC suppresses inflammatory responses in macrophages.

## 1. Introduction

Immunosuppressive effects of environmental toxicants are an important but poorly understood mechanism of toxicity (Hartung and Corsini 2013). Environmental toxicant suppression of immune responses potentially leads to increased susceptibility to infectious diseases in exposed populations, thereby contributing to the overall public health burden of pathogenic organisms such as viruses and bacteria (Repetto and Baliga 1997, Birnbaum and Jung 2010, Feingold, Vegosen et al. 2010, Hartung and Corsini 2013, Anderson and Shane 2018). Studies in humans and animal models have identified potential immunosuppressive effects of environmental toxicants. For example, children exposed to perfluorinated compounds have decreased antibody response to vaccines (Grandjean, Andersen et al. 2012, Granum, Haug et al. 2013) and mice chronically exposed to arsenic have decreased expression of cytokines (Yan, Xu et al. 2020). However, many environmental toxicants have not been examined for immunosuppressive hazards.

Trichloroethylene is an organic solvent used in industrial applications as a metal degreaser, in the manufacturing of refrigerants and in dry cleaning operations (ATSDR 2019). Trichloroethylene is found in hundreds of Superfund sites and is a frequent groundwater contaminant (Chiu, Jinot et al. 2013, Makris, Scott et al. 2016). Exposure can occur via contaminated drinking water or air near waste-sites or factories that use trichloroethylene (ATSDR 2019). Trichloroethylene is an immunotoxicant (Aranyi, O’Shea et al. 1986, Zhang, Bassig et al. 2013, Zhang, Li et al. 2017), as shown in studies in which trichloroethylene-metabolizing enzymes were inhibited suggest that downstream metabolites are responsible for at least some of trichloroethylene’s immunotoxic effects, as opposed to the parent compound (Griffin, Gilbert et al. 2000, Lash, Chiu et al. 2014). We previously demonstrated that S-(1,2-dichlorovinyl)-l-cysteine (DCVC), a bioactive trichloroethylene metabolite, can suppress the immune/inflammatory response to a common pathogen (group B *streptococcus*) in fetal membrane tissue *ex vivo* (Boldenow, Hassan et al. 2015). These immunosuppressive effects included decreases in pathogen-stimulated secretion of several important cytokines such as TNF-α, IL-1β and IL-8. Similar effects were also observed when inflammatory responses were induced by cell wall components from gram-positive (lipoteichoic acid) and gram-negative bacteria (lipopolysaccharide). However, the *ex vivo* fetal membrane model used in this study were made using whole-tissue punches, which contain an array of cell types including fibroblasts, trophoblasts, epithelial cells and resident macrophages (Uchide, Ohyama et al. 2012). Thus, how DCVC affects specific immune cell types and the mechanisms involved, is still not well understood.

To more fully understand how DCVC impacts on immune processes, we used next generation sequencing (RNA-sequencing) to identify transcriptomic pathways impacted by DCVC under conditions of immune stimulation in macrophages, a key immune cell type found in various tissues throughout the body including the fetal membranes. Understanding DCVC impacts on macrophage gene expression and cytokine levels is critical to understanding the immunosuppressive effects of TCE exposure, given the previously observed immunosuppressive effects observed in the fetal membranes. Impacts on macrophages could suggest effects in other tissues and organs, providing important context to the health hazards posed by trichloroethylene exposure. In this study, we paid a particular focus on the Toll-Like Receptor 4 (TLR4) signaling pathway due to its important role in stimulating inflammatory responses in immune cells (Palsson-McDermott and O’Neill 2004).

## 2. Materials and Methods

### 2.1 Cell culture

THP-1 cells were obtained from American Type Culture Collection. THP-1 cells were selected because they are a relevant human monocyte cell line that can be differentiated into a macrophage like cell that responds similarly to placental macrophages (Tetz, Aronoff et al. 2015, Tsai, Tseng et al. 2017). THP-1 cells were cultured using standard procedures at 37°C and 5% CO_2_. The cell culture media was RPMI 1640 supplemented with 10% FBS and penicillin/streptomycin (pen/strep). Twenty-four hours prior to treatment, cells were plated in 24-well plates at a concentration of approximately 2×10^5^ cells/mL with 100 nM phorbol 12-myristate 13-acetone (PMA). Following differentiation with PMA, cells were treated with medium only or DCVC (1, 5, or 10 μM) for another 24 hours. DCVC was synthesized by the University of Michigan Medicinal Chemistry Core Synthesis Lab (purity >98%) (Boldenow, Hassan et al. 2015, Hassan, Kumar et al. 2016). The DCVC concentrations were selected based on the concentration range of detection of trichloroethylene metabolites in human serum and we have previously observed impacts in human gestational tissues at these levels (Lash, Putt et al. 1999, Boldenow, Hassan et al. 2015, Hassan, Kumar et al. 2016, ATSDR 2019). After DCVC treatment, medium was changed and lipopolysaccharide (LPS; 100 ng/mL) was added to selected wells for 4 h. Treatment conditions included: medium only (control), LPS (100 ng/mL), DCVC (1, 5, or 10μM), and DCVC (1, 5, or 10μM) + LPS (100 ng/mL). Media was collected for LDH cytotoxicity and cytokine analysis and RNA was collected for RNA-seq. LDH and cytokine experiments were conducted three times (n=3) in triplicate.

### 2.2 LDH Cytotoxicity Assay

Cytotoxicity across all treatment groups was assays using Takara (Kusatsu, Shiga, Japan) lactate dehydrogenase (LDH) cytotoxicity detection kit and following kit protocols. Briefly, after treatment with medium only (control), LPS (100 ng/mL), DCVC (1, 5, or 10μM), and DCVC (1, 5, or 10μM) + LPS (100 ng/mL), cell culture media was collected and LDH activity was measured by addition of a color changing substrate followed by quantification of 490/492 absorbance using a BioTek (Winooski, VT) Eon™ Microplate Spectrophotometer plate reader.

### 2.3 RNA Sequencing

RNA was extracted from THP-1 cells for the following four treatment groups: control (no treatment), LPS treatment (100ng/mL), DCVC (5µM) treatment or LPS (100ng/mL) + DCVC (5µM) co-treatment (n=6 per treatment group). RNA was extracted using the Quick-RNA MiniPrep kit and standard protocol (Zymo Research, Irvine, CA). RNA was stored at -80°C. The University of Michigan Advanced Genomics Core performed RNA sequencing. RNA library preparation was performed using poly-A selection with TruSeq Stranded mRNA Library Prep Kit (lllumina; San Diego, CA, USA) on the NovaSeq-6000 platform (Illumina). RNA library preparation was performed using poly-A selection. Paired end sequencing with 150bp sequences was done on the NovaSeq-6000 platform (Illumina).

### 2.3 RNA-seq Processing

Raw fastq files were first examined using fastQC (version 0.11.5) (Andrews 2010), and reports generated for all 24 samples (six sample per treatment group*four treatment groups) were collated using multiQC (version 0.9) (Ewels, Magnusson et al. 2016). Mean quality scores across all base positions were high for all samples. Reads were mapped to the human reference genome (hg38) using the Spliced Transcripts Alignment to a Reference (STAR) (version 2.6.0c) program (Dobin, Davis et al. 2013). QoRTs (version 1.3.6) (Hartley and Mullikin 2015) was used to examine post alignment quality control metrics. Approximately 3-6% of reads were dropped in each sample for multi mapping, and approximately 90% of read mapping locations were unique gene regions. Following mapping, featureCounts (version 1.6.1) (Liao, Smyth et al. 2014) was used to quantify aligned reads mapping to exons.

### 2.4 Differential Gene Expression and Principal Components Analysis (PCA)

Following alignment and quantification, data were evaluated for differential gene expression. Gene counts were read into R (version 3.6.0), and analyzed with the DESeq2 package (version 1.24.0) (Love, Huber et al. 2014). Principal components were plotted, calculated on variance stabilizing transformed values of the expression data, to examine clustering. Principal component calculation used the top 500 genes ranked by variance. Data were evaluated for differential gene expression by treatment group. In DESeq2, model terms were treatment group (four levels: control, DCVC, LPS and LPS+DCVC). Initially each treatment group (DCVC, LPS and LPS+DCVC) was compared to the control group. For the purposes of this study, DCVC alteration of the inflammation response to LPS was of particular interest, therefore an additional comparison was made between LPS+DCVC treated cells vs. LPS treated cells. A Benjamini-Hochberg adjusted p-value < 0.05 and an absolute log_2_fold-change > 2.5 threshold to determine significance was used.

Default settings for DESeq2 were used for filtering of genes with low normalized mean counts. Volcano plots of results using the EnhancedVolcano (version 1.2.0) package were created after applying log fold-change shrinkage using the “apeglm” (Zhu, Ibrahim et al. 2019).

### 2.5 Gene Set Enrichment Analysis

We tested differentially expressed genes for enrichment of gene ontology terms using Gene Set Enrichment Analysis (GSEA) (Ashburner, Ball et al. 2000, Subramanian, Tamayo et al. 2005). Results from DESeq2 sorted by the values for the Wald test statistic were used as input to test for enrichment against ontology gene sets (Ashburner, Ball et al. 2000) downloaded from the Molecular Signatures Database (v7.4) (Liberzon, Subramanian et al. 2011). DCVC vs control, LPS vs control, DCVC+LPS vs control and DCVC+LPS vs LPS were the comparisons of interest for GSEA. Gene ontology terms with adjusted p-value<0.05 were significantly enriched and we reported normalized enrichment scores (NES).

### 2.6 Toll-like receptor 4 (TLR4) Signaling Pathway

The toll-like receptor 4 (TLR4) signaling pathway was adapted from multiple sources (Guha and Mackman 2001, Palsson-McDermott and O’Neill 2004). We visualized relative changes in gene expression in this pathway for LPS+DCVC co-treated samples vs. samples treated with LPS alone using a network diagram. All genes in the pathway were plotted without a significance threshold for inclusion.

### 2.7 RNA-seq Data Availability

Raw and processed data can be accessed at the genome expression omnibus (accession number pending). Code for analysis is available (https://github.com/bakulskilab).

### 2.8 Cytokine Quantification

Cytokine concentrations in cell culture media were quantified for the following treatments: 1) control (no treatment), LPS (100ng/mL), DCVC alone (1, 5 or 10µM) or DCVC (1, 5 or 10µM) + LPS (100ng/mL) co-treatments (n=3 per treatment group). Cytokines were quantified using ELISA kits from R&D Systems (Minneapolis, MN) and following kit protocols. Cytokines quantified included TNF-α, IL-6, IL-1β. Detection ranges by cytokine were 15.6-1,000 pg/mL for TNF-α, 9.38-600 pg/mL for IL-6, and 3.9 - 250 pg/mL for IL-1β. Samples below the limit of detection (LOD) were transformed to LOD/√2 prior to statistical analysis.

### 2.9 Statistical Analysis for Cytokine Levels

Values were averaged for replicated within each experiment. Data are presented as mean ± SEM and were analyzed using GraphPad Prism software (GraphPad Software, La Jolla, CA). ANOVAs with Tukey’s post hoc test were performed. Data were considered significant if the p-value was < 0.05.

## 3. Results

### 3.1 LDH Cytotoxicity Assay

DCVC (1, 5, and 10 µM), LPS, and DCVC (1, 5, and 10 µM) + LPS treatments did not alter cell death compared to control as measured by LDH release (data not shown).

### 3.2 Principal components analysis of transcriptomic profiles

Principal components analysis of transcriptomic profiles showed a clear separation for all the treatment groups (control, DCVC treated, LPS treated, DCVC+LPS treated) (Figure 1). Principal component 1 captured 78% of the variance and showed a clear separation between samples treated with LPS and samples not treated with LPS. Principal component 2 captured 16% of the variance and showed a clear separation between samples treated with DCVC and samples not treated with DCVC (Figure 1).

**Figure. 1.**
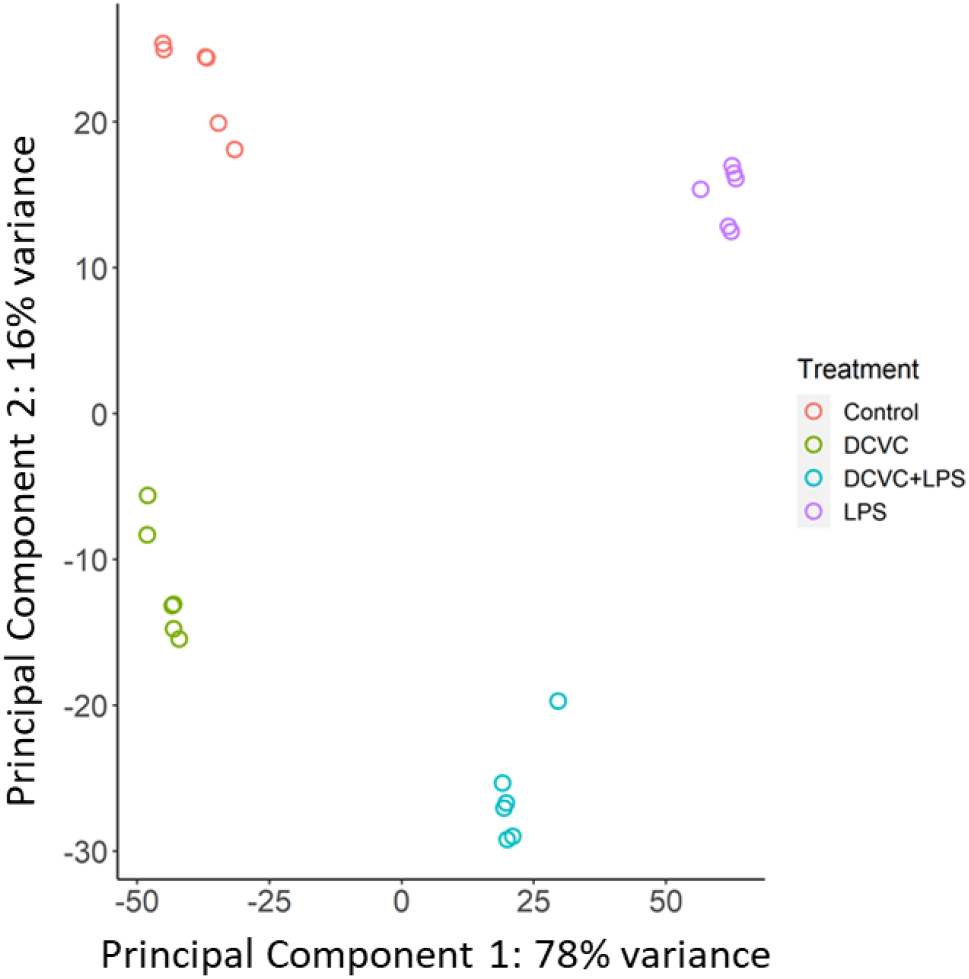
Principal components analysis of global gene expression for all treatment groups. THP-1 cells were cultured and treated with one of the following experimental conditions: control (no treatment), LPS (100ng/mL), DCVC (5µM) or LPS (100ng/mL) +DCVC (5µM). Total RNA from each was isolated and transcriptomic profiles were generated using RNA-seq. Transcriptomic profiles were then grouped by similarity using principal components analysis.

### 3.3 Significant Genes Changed Across Treatment Groups

There were 826 differentially expressed genes between DCVC treated cells versus controls, as defined by our statistical significance criteria (adjusted p-value < 0.05 and log_2_fold-change magnitude > 2.5). Of the differentially expressed genes, 506 genes were upregulated and 320 were downregulated (Figure 2A). For example, *PIR* was 6-fold upregulated with DCVC treatment (adjusted p-value=9.7*10^−277^) and *TNFAIP8L3* was 12-fold downregulated (adjusted p-value=8.4*10^−57^). The top 10 significantly changed genes (ranked by adjusted p-value) were: *PIR, GCLM, IGGA1, MYOF, BCL2A1, UST, FAM49A, LRRC25, RGS16* and *OSGIN1*. A full list of differentially expressed genes for DCVC vs. control treatments is shown in Supplemental Table 1.

**Figure. 2.**
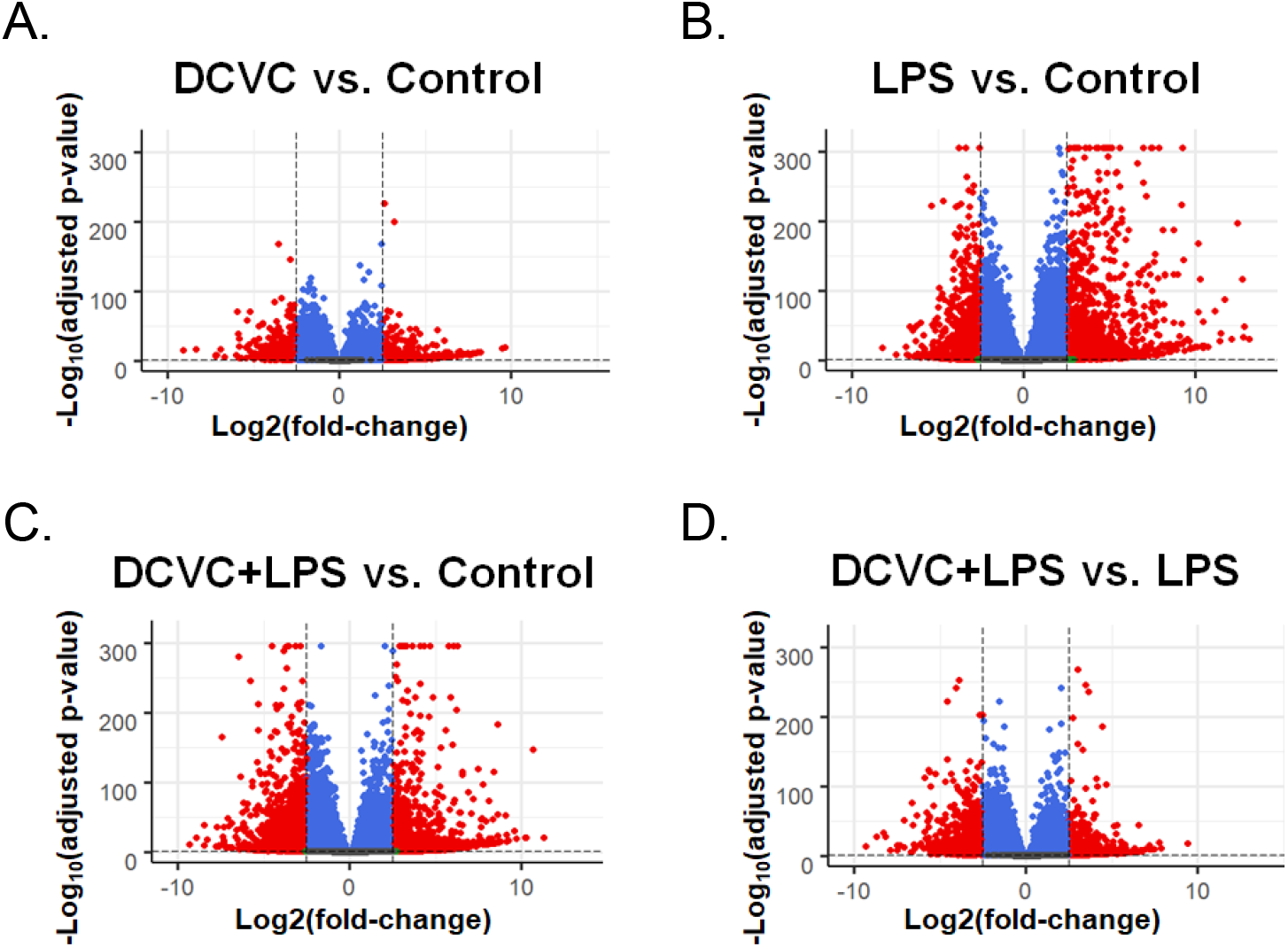
Volcano plots showing significant gene changes across DCVC and/or LPS treated THP-1 cells. Global gene expression profiles across all treatment groups were analyzed using RNA-seq. Volcano plots show log_2_fold-changes and – log_10_P-values for comparisons between DCVC vs. Control (A), LPS vs. Control (B), DCVC+LPS vs. Control (C) and DCVC+LPS vs. LPS (D) treatments. Significantly changed genes (adjusted p-value < 0.05 and an absolute log_2_fold-change > 2.5) are shown as red dots and non-significantly changed genes are shown as blue dots.

When comparing LPS treatment versus controls, 1,989 genes were differentially expressed (1,107 upregulated, 882 downregulated, Figure 2B). For example, *TNFAIP8L3* was 40-fold upregulated with LPS treatment (adjusted p-value=1.0*10^−269^) and *PLA2G15* was 7-fold downregulated (adjusted p-value=2.1*10^−251^). The top 10 significantly changed genes (ranked by p-value) were: *SOCS3, TRAF1, MCOLN2, TNFAIP2, ABTB2, CD83, NFKBIZ, KYNU, XIRP1* and *REL*. A full list of differentially expressed genes for LPS vs. control treatments is shown in Supplemental Table 2.

When comparing LPS+DCVC co-treatment versus controls, 2,055 genes were differentially expressed (1,100 upregulated, 955 downregulated, Figure 2C). For example, *NFKB1* was 7-fold upregulated with LPS+DCVC treatment (adjusted p-value=6.0*10^−269^) and *ST3GAL5* was 15-fold downregulated (adjusted p-value=1.8*10^−235^). The top 10 significantly changed genes (ranked by p-value) were: *TRAF1, TNFAIP2, ABTB2, GCLM, XIRP1, TXNRD1, TNIP1, RFFL, PIR* and *WTAP*. A full list of differentially expressed genes for LPS vs. control treatments is shown in Supplemental Table 3.

Finally, 1,399 genes were differentially expressed in cells co-treated with LPS+DCVC vs. LPS treatments (773 upregulated, 626 downregulated, Figure 2D). For example, *GCLM* was 11-fold upregulated with LPS+DCVC treatment (adjusted p-value=1.8*10^−245^), *MYOF* was 15-fold downregulated (adjusted p-value=1.3*10^−252^) and *IDO1* was 56-fold downregulated (adjusted p-value=7*10^−26^). The top 10 significantly changed genes (ranked by p-value) were: *PIR, MYOF, GCLM, LDLRAD3, PLEKHM3, RGS16, B4GALT5, STK17B, GSR* and *OSGIN1*. A full list of differentially expressed genes for LPS+DCVC vs. LPS treatments is shown in Supplemental Table 4.

### 3.4 Geneset Enrichment Analysis: DCVC, LPS or DCVC+LPS Co-treatments vs. Controls

After removing redundant terms, geneset enrichment analysis revealed 580 significantly enriched biological pathways (i.e. Gene Ontology defined biological processes) for DCVC treated cells, 365 enriched pathways in LPS treated cells and 359 enriched pathways in DCVC+LPS co-treated cells. The full lists of Gene Ontology terms enriched for each treatment group are shown in Supplementary Tables 5-8. Figure 3 shows the number of common versus uniquely enriched processes across all three treatment groups. As shown in Figure 3, there were 51 processes enriched across all three treatments. Compared to the control treatment group, there were 228 significantly enriched processes in DCVC+LPS co-treatments which were not enriched with LPS treatment alone (see Figure 3), indicating that DCVC caused a robust alteration in the transcriptomic response to LPS stimulation.

**Figure. 3.**
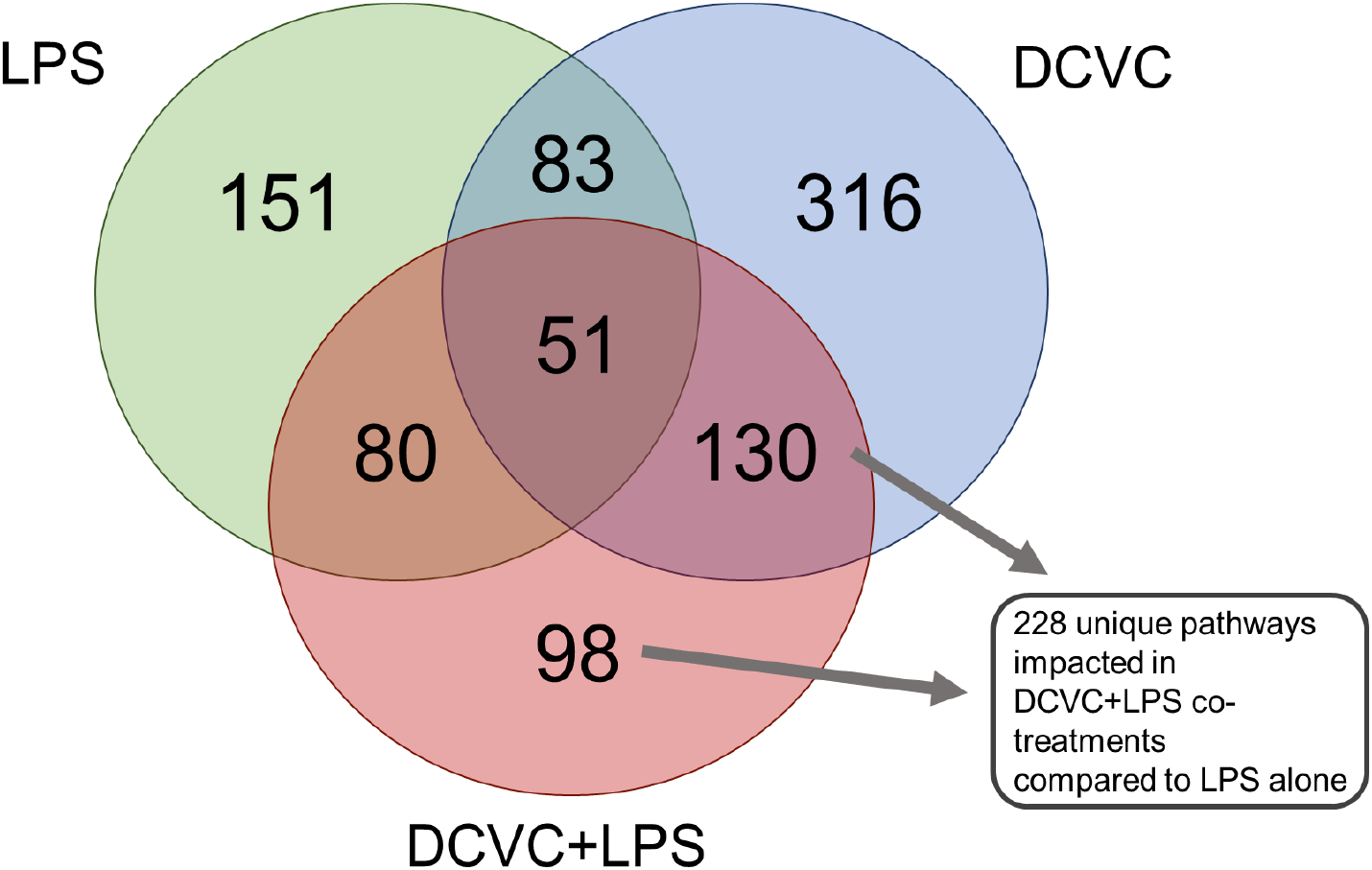
Number of overlapping or unique significantly enriched biological processes across all treatment groups. After treatment, RNA was isolated from THP-1 cells and transcriptomic profiles were analyzed using RNA-seq. Venn diagrams show the number of Biological Processes commonly or uniquely enriched across the three treatment groups compared to the untreated control group. We observed that LPS+DCVC co-treatments were associated with 228 enriched Biological Processes that were not enriched after treatment with LPS alone.

### 3.5 Gene Set enrichment analysis: DCVC+LPS co-treatments vs. LPS-only treatments

When directly comparing DCVC+LPS co-treatment to LPS-only treatment groups, we identified 626 enriched biologic processes after removing redundant terms. The top ten pathways (ranked by adjusted p-value) are shown in Table 1. Pathways with a negative enrichment score (i.e. pathways overrepresented among genes downregulated in DCVC+LPS vs. LPS-only treatments) included several pathways related to immune cell activation (“leukocyte activation”, adjusted p-value=5.4*10^−20^, NES=-2.2), cell signaling pathways (“positive regulation of MAPK cascade”, adjusted p-value=1.0*10^−21^, NES=-2.2) and cytokine production (“positive regulation of cytokine production”, adjusted p-value=8.4*10^−19^, NES=-2.2). Additional downregulated immune/inflammation pathways included “acute inflammatory response” (adjusted p-value=3.5*10^−5^, NES=-1.9), “production of molecular mediator of immune response” (adjusted p-value=3.1*10^−7^, NES=-2.0) and “response to virus” (adjusted p-value=2.9*10^−18^, NES=-2.3) (see Supplementary Table 8). As shown in Table 2 significantly impacted immune/inflammation pathways also included multiple interleukin (IL) pathways such as “IL-7 mediated signaling pathway” (adjusted p-value=0.01, NES=-1.7), “response to interleukin-12” (adjusted p-value=2.4*10^−5^, NES=-2.1), “response to interleukin-15” (adjusted p-value=0.0006, NES=-1.9) and “response to interleukin-18” (adjusted p-value=0.005, NES=-1.8) as well as pathways involved in macrophage functions, including “phagocytosis” (adjusted p-value=6.3*10^−7^, NES=-1.8) and “cell killing” (adjusted p-value=0.001, NES=-1.7). In contrast, upregulated pathways (i.e. pathways overrepresented among genes downregulated in DCVC+LPS vs. LPS-only treatments) included those related to xenobiotic metabolism, such as “glutathione metabolic process” (adjusted p-value=0.005, NES=1.7), “xenobiotic catabolic process” (adjusted p-value=0.04, NES=1.6) and “detoxification” (adjusted p-value=0.02, NES=1.5).

**Table 1.** Top 10 significantly enriched pathways for LPS+DCVC vs. LPS-only treatment groups.

| Biological Process<br>(Gene Ontology ID#) | Adjusted p-value | Normalized Enrichment<br>Score |
| --- | --- | --- |
| Regulation of cell activation<br>(GO:0050865) | $4 \times 10^{-26}$ | -2.3 |
| Ribonucleoprotein complex<br>biogenesis (GO:0022613) | $1 \times 10^{-23}$ | -2.2 |
| Positive regulation of MAPK cascade<br>(GO:0043410) | $1 \times 10^{-21}$ | -2.2 |
| Positive regulation of locomotion<br>(GO:0040017) | $2 \times 10^{-22}$ | -2.2 |
| Regulation of lymphocyte activation<br>(GO:0051249) | $5.4 \times 10^{-20}$ | -2.2 |
| Cellular response to lipid<br>(GO:0071396) | $5.4 \times 10^{-19}$ | -2.1 |
| Positive regulation of cytokine<br>production (GO:0001819) | $8.4 \times 10^{-19}$ | -2.2 |
| Peptidyl tyrosine modification<br>(GO:0018212) | $2.4 \times 10^{-18}$ | -2.3 |
| Response to virus (GO:0009615) | $2.9 \times 10^{-18}$ | -2.3 |
| Positive regulation of protein kinase<br>activity (GO:0045860) | $3.2 \times 10^{-18}$ | -2.1 |

**Table 2.**
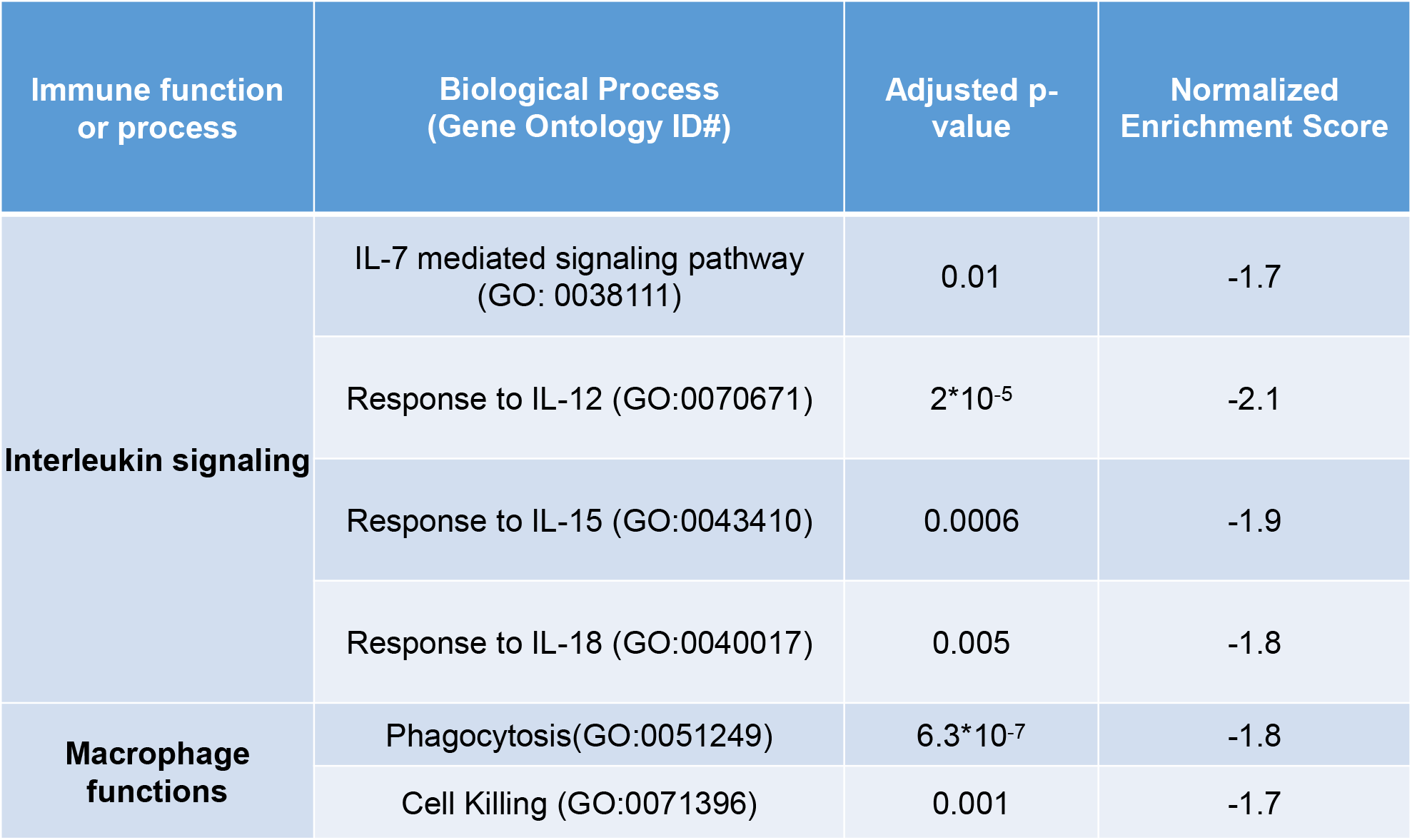
Significantly downregulated immune pathways in LPS+DCVC vs. LPS-only treatment groups.

### 3.6 Toll-like receptor 4 (TLR4) Signaling Pathway

Multiple genes were downregulated in the TLR4’s cellular pathway in DCVC+LPS co-treatment compared to the LPS treatment group (see Figure 4). Genes coding for proinflammatory cytokines (for example, TNF-α and IL-6) and the cell adhesion molecule *VCAM1* (Kong, Kim et al. 2018) showed the largest degree of downregulation (log_2_fold-change ≤ -3.5; adjusted p-value < 0.001). Genes coding for *TLR4, BTK, MYD88, IRAK-1, NF-*κ*B*, and *ELK-1* were also downregulated (log_2_fold-change ≤ -0.7; adjusted p-value ≤ 0.001). Conversely, DCVC+LPS co-treatment relative to LPS alone upregulated LBP and cFos (log_2_fold-change ≥ 1.0; adjusted p-value < 0.05).

**Figure. 4.**
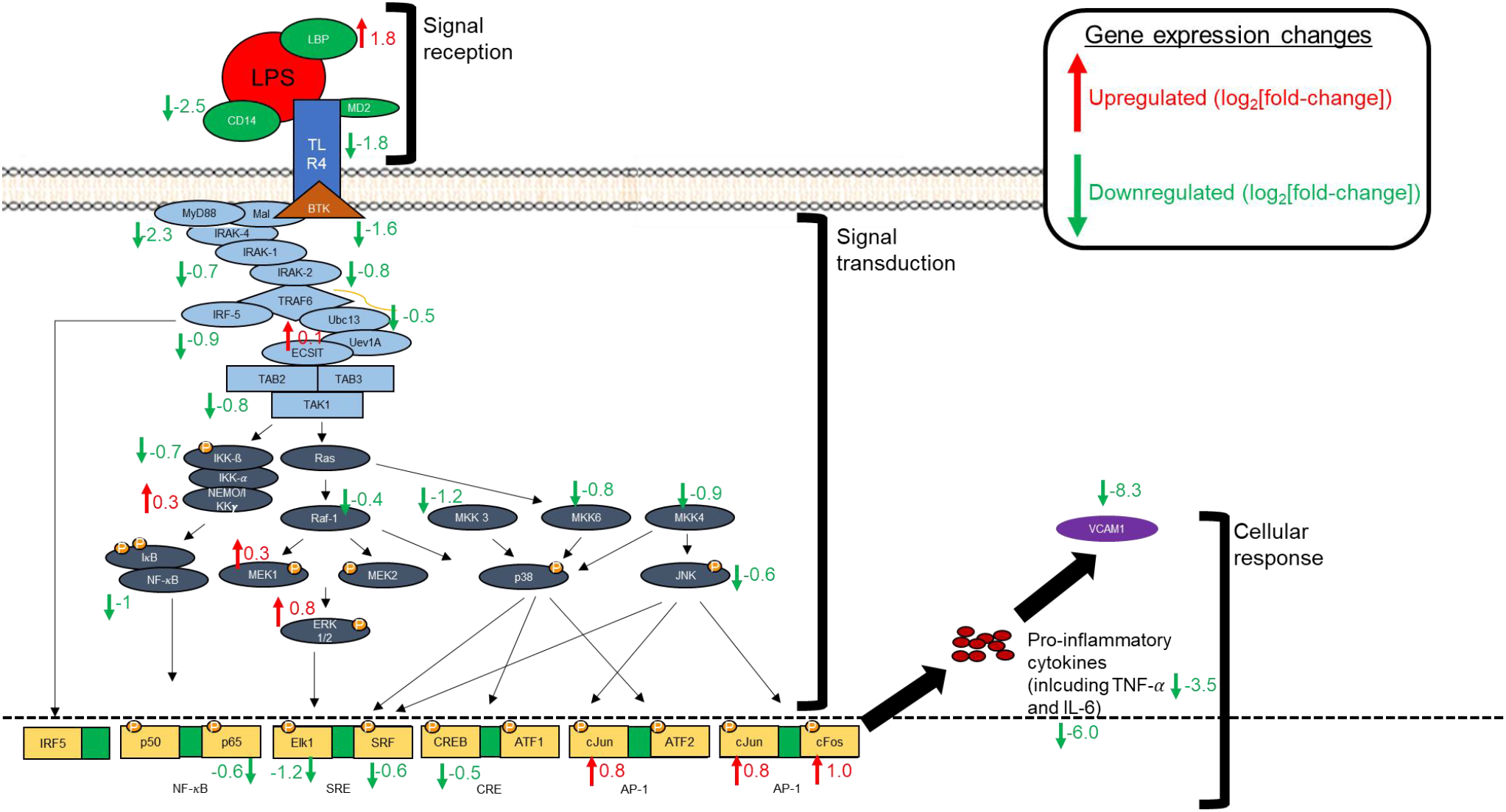
Gene expression changes for genes in the Toll-like receptor 4 (TLR4) Signaling Pathway. The Toll-like receptor 4 (TLR4) pathways was adapted using previously published sources (Guha and Mackman 2001, Palsson-McDermott and O’Neill 2004). Gene expression changes for DCVC+LPS vs. LPS-only treatments are shown for genes in the pathway. Most of the genes in the pathway were downregulated in DCVC+LPS co-treatments compared to LPS-only treatments.

### 3.7 Cytokine analysis

Co-treatment of THP-1 cells with DCVC (1, 5, or 10 μM) and LPS (100 ng/mL) for 4 hours, decreased cytokine levels in a dose dependent manner (Figure 5). DCVC treatment alone did not alter cytokine release relative to controls. LPS treatment alone caused an approximately 118-fold increase in TNF-α release, a 24-fold increase in IL-6 release, and a 52-fold increase in IL-1β release compared to controls (p < 0.05). Co-treatment of DCVC and LPS caused a substantial dose-dependent decrease in levels of cytokine release 4 hours post-treatment. Comparing DCVC+LPS co-treatment to LPS treatment, TNF-α showed a 1.7-fold decrease at 1 μM DCVC, a 7-fold decrease at 5 μM DCVC, and a 22-fold decrease at 10 μM DCVC (add p-values). In the DCVC+LPS co-treatment, compared to the LPS treatment, IL-6 showed a 6-fold decrease at 1 μM DCVC, and no measurable release at 5 or 10 μM DCVC, or an approximate 24-fold decrease in IL-6 release. DCVC inhibition of LPS stimulated IL-6 was statistically different at all three DCVC concentrations (1, 5 and 10 µM). With LPS + DCVC IL-1β showed an 8-fold decrease at 1 μM, a 20-fold decrease at 5 μM, and a 27-fold decrease at 10 μM compared to the LPS treatment. DCVC inhibition of LPS stimulated IL-1β was statistically different at all three DCVC concentrations (1, 5 and 10 µM).

**Figure. 5.**
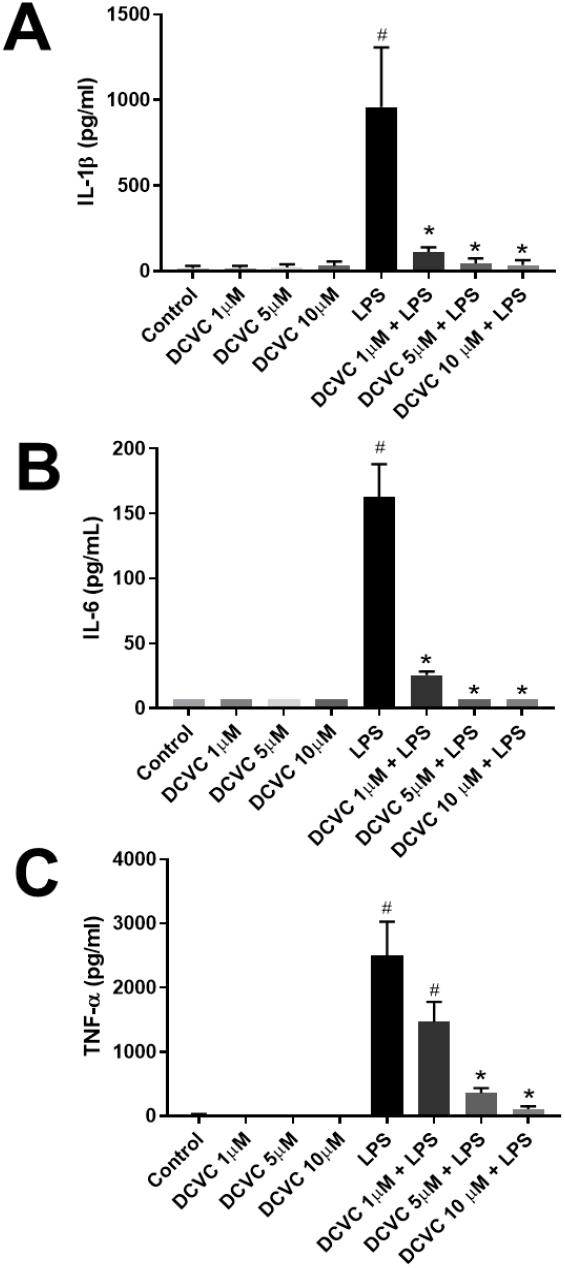
DCVC effects on LPS-stimulated cytokine secretion. DCVC effects on LPS-stimulated release of pro-inflammatory cytokines from THP-1 cells. THP-1 cells were differentiated with PMA and then treated with DCVC for 24 h. Following DCVC treatment, cells were treated with LPS for 4 h and then medium was collected and assayed by ELISA for IL-1β (A), IL-6 (B), and TNF-α (C). Columns represent mean ± SEM. Data were analyzed by ANOVA with Tukey’s post-hoc test. #, Significant differences compared to control (medium only) (p≤0.05). *, Significant differences compared to LPS alone (p≤0.05).

## 4. Discussion

Trichloroethylene contamination remains a public health concern due to large scale production and release into the air, drinking water, and soil (ATSDR 2019). In a recent risk evaluation of trichloroethylene, the United States Environmental Protection Agency (USEPA) found “unreasonable risks” to human health for consumers using products containing trichloroethylene (e.g. cleaning products, arts and crafts spray coatings) as well as workers using trichloroethylene directly or those working nearby (occupational non-users) (USEPA 2020). As part of this risk evaluation, the USEPA noted immunotoxic effects that included autoimmunity/inappropriate immune activation as well as immunosuppression and cited these endpoints as the most important non-cancer endpoint for risk conclusions (USEPA 2020). Importantly, gaps still remain in our understanding of molecular pathways that underpin the immunosuppressive effects of trichloroethylene.

In this study, we sought to identify molecular pathways impacted by the bioactive trichloroethylene metabolite, DCVC, in macrophages after activation of inflammation pathways. We demonstrated that macrophages are potentially a key target of the immunosuppressive activity of DCVC. This study provides cell type specific molecular evidence supporting earlier studies in our lab showing that DCVC suppresses immune responses in fetal membrane tissue (Boldenow, Hassan et al. 2015). Moreover, because macrophages are found in virtually every major organ (Ovchinnikov 2008), these finding have important implications for immunosuppression in multiple tissue and organ systems.

Our findings show that the trichloroethylene metabolite DCVC strongly inhibits key transcriptional pathways related to inflammatory responses and macrophage functions in a macrophage cell model. Pathways inhibited included multiple interleukin pathways which play important roles in mediating inflammation and immune responses to pathogens. These included IL-6 (Tanaka, Narazaki et al. 2014), IL-7 (Willis, Seamons et al. 2012), IL-12 (Trinchieri 2003), IL-15 (Perera, Lichy et al. 2012), IL-18 (Nakanishi 2018), IL-27 (Iwasaki, Fujio et al. 2015) and IL-35 (Sawant, Hamilton et al. 2015) as well as the TLR4 pathway (Molteni, Gemma et al. 2016). Moreover, DCVC inhibits the release of important pro-inflammatory cytokine proteins including TNF-α, IL-6 and IL-1β. Our transcriptional analysis of the TLR4 pathway suggests that DCVC suppression these cytokines occurs upstream of cytokine protein translation, e.g. via suppression of key genes such *TLR4* or *CD14*. Additional pathways related to immune functions or inflammation (for example: “acute inflammatory response”, “positive regulation of cytokine production” and “regulation of lymphocyte activation”) were significantly downregulated in DCVC+LPS treatments versus LPS-only treatments, suggesting that DCVC downregulates critical LPS-induced immune and inflammatory pathways in macrophages. Importantly downregulated pathways included those directly related to macrophage immune function such as “phagocytosis” and “cell killing”. Finally, the gene for indoleamine 2,3-dioxygenase 1 (*IDO1*), was significantly downregulated by DCVC treatment (55-fold lower expression in LPS+DCVC co-treatments compared to LPS-only treatments). Indoleamine 2,3-dioxygenase 1 is a metabolic enzyme that converts tryptophan to it’s metabolite, kynurenine (Hornyak, Dobos et al. 2018). IDO1 has been shown to have a multiple roles in regulating immune responses, e.g. in immune tolerance and in limiting tissue damage associated with inflammation (Munn and Mellor 2016). DCVC suppression of IDO1 and the role this effect plays in the immunomodulatory effects of TCE exposure warrants further study. In contrast to downregulated pathways, pathways upregulated in the DCVC + LPS treatments compared to LPS-only treatments such as “glutathione metabolic processing”, “detoxification” and “xenobiotic metabolic processing” were likely due to cellular metabolism of DCVC, as DCVC is a product of the glutathione conjugation pathway of trichloroethylene metabolism and is itself metabolized by enzymes in this pathway (Lash, Fisher et al. 2000). Taken together our transcriptomic analysis provides important information about how DCVC alters the inflammatory response in macrophages.

Consistent with what we previously observed previously in fetal membranes, our cytokine analysis showed that the DCVC and LPS co-treatments significantly suppressed LPS-stimulated release of proinflammatory cytokines such as TNF-α, IL-6, and IL-1β compared to treatment with LPS alone (Boldenow, Hassan et al. 2015). TNF-α, one of the most important proinflammatory cytokines, has many roles in the innate immune system functioning, including vasodilatation, edema formation, leukocyte adhesion to epithelium and regulation of blood coagulation. TNF-α also contributes to oxidative stress at sites of inflammation (Semenzato 1990) and is also vital for the recruitment of neutrophils and the promotion of macrophage phagocytosis (Lukacs, Strieter et al. 1995, Arcuri, Toti et al. 2009). Therefore, downregulation of TNF-α could have significant and wide-ranging implications on the body’s immune system functioning and cells’ susceptibility to infection.

IL-6 is another important proinflammatory cytokine that has been implicated in the body’s immune response inflammation, and hematopoiesis. It is released in significant amount by monocytes and macrophages during infection to assist and activate the immune system for proper clearance of pathogens (Tanaka, Narazaki et al. 2016). Overproduction of IL-6, on one hand, has been implicated in autoimmune disorders and chronic inflammation, and IL-6 blockades have been found to be successful in treating them (Tanaka, Narazaki et al. 2014). However, the most common side effect is an increased susceptibility to severe infections (Choy, De Benedetti et al. 2020). Therefore, DCVC suppression of IL-6 may have the effect of increasing a cell’s risk of infection.

The pro-inflammatory cytokine IL-1β is secreted in an inactive form before being cleaved and activated by a caspase cascade (Yazdi and Ghoreschi 2016). It is not secreted typically through the endosome, leading some researchers to suggest that its method of production varies depending on the cell’s needs (Lopez-Castejon and Brough 2011). Because of the complex nature by which IL-1β is secreted and its numerous and often indirect on inflammation, more research is needed on the full implications of DCVC suppression of IL-1β secretion.

The concentrations of DCVC chosen for this study were based on known bioactive concentrations in kidney cells and human placental cells (Xu, Papanayotou et al. 2008, Hassan, Kumar et al. 2016, Elkin, Harris et al. 2018). Although blood levels of DCVC in trichloroethylene exposure scenarios can be difficult to verify, humans exposed to the current occupational exposure limit of 100 ppm trichloroethylene in air can have DCVG (a precursor to DCVC) serum levels as high as 50 μM (Lash, Putt et al. 1999). Typical environmental concentrations of trichloroethylene are <5 ppb in air (ATSDR 2019, USEPA 2020). Therefore, although the concentrations of DCVC in the present study were higher than the average ambient environmental exposure in the US, they are within plausible limits found in occupational exposures.

Despite the paradigm shift underway within toxicology to focus more on interactions of multiple factors, there remains scare knowledge about the implications of pathogen-toxicant interactions for human health. Here, we showed that DCVC suppresses the macrophage inflammation response in response to LPS at both the gene expression and cytokine level. This work has broad implications because macrophages are found throughout the body and participate in many important roles that include maintaining homeostasis, responding to infections, resolving inflammation and repairing tissue damage (Mosser and Edwards 2008, Murray and Wynn 2011). Because cytokine secretion being is the foundation of immune cell recruitment, DCVC inhibition of macrophage responses to pathogens could result in decreased recruitment of immune cells and potentially increase the number and severity of pathogenic infections in individuals exposed to trichloroethylene.

It is important to note that the experiments of the present study were done using a macrophage cell line *in vitro*. Moreover, macrophages are also highly heterogenous cells that adapt quickly to their microenvironments, and help direct inflammation responses in both proinflammatory and anti-inflammatory directions (Mosser and Edwards 2008, Murray and Wynn 2011). Because of their adaptability, particularly regarding their ability to respond to and directly modify the immune system’s inflammation response, future studies in physiologically relevant animal models and in primary macrophages obtained from multiple tissues are needed to better understand DCVC’s immunomodulatory effects. For example, to confirm the results observed in this study and more extensively assess the pathways and macrophage functions impacted by DCVC, further experiments are needed to verify responses observed in the THP-1 cell line using primary macrophages (e.g. human CD14+ cells isolated from peripheral blood mononuclear cells) and assess impacts on cytokines using full cytokine/chemokine arrays. In addition, future experiments should investigate DCVC impacts on macrophage functions such as phagocytosis and cell killing activity. Finally, *in vivo* models could help to confirm the full integrated immune response to DCVC at the organism level.

## 5. Conclusion

Our results demonstrate that DCVC suppresses multiple LPS-stimulated transcriptional inflammation pathways and expression of pro-inflammatory cytokines in macrophages. Suppression of inflammatory pathways occurred upstream of cytokine protein expression and included multiple interleukin pathways as well as the TLR4 pathway. Our results suggest that TCE exposure could lead to suppression of key macrophage responses to pathogenic infection via its bioactive metabolite, DCVC.

## Acknowledgements

This work was supported by Michigan Center on Lifestage Environmental Exposures and Disease (P30ES017885). Drs. Harris, Bakulski and Loch-Caruso were supported by the National Institute of Environmental Health Sciences of the National Institutes of Health under Award Number P42ES017198. Additional funding for Dr. Harris was provided by the National Center for Advancing Translational Sciences (UL1TR002240). Dr. Bakulski was supported by grants from the National Institutes of Health (R01ES025531; R01ES025574; R01AG055406; R01MD013299).

